# Investigating the antibacterial and anti-biofilm activity of agave syrup as a traditional wound remedy

**DOI:** 10.1101/2025.07.13.664611

**Authors:** Ross Pallett, Kathleen Pritchard, Kayleigh Wilkins, Raphael Galleh, Olusoji Adebisi, Lucy Ann Harman, Talvin Momi, Harvey Wilkes, Caroline Dodds Pennock

**Affiliations:** School of Biosciences, College of Health and Life Sciences, Aston University, Birmingham, United Kingdom; School of History, Philosophy and Digital Humanities, University of Sheffield, Sheffield, United Kingdom

**Author notes:** **Corresponding author:** Ross Pallett School of Biosciences, College of Health and Life Sciences, Aston University, Birmingham, United Kingdom. Raphael Galleh - Institute of Science and Environmental Science, University of Cumbria, Cumbria, United Kingdom. These authors contributed equally to this work (Lucy Ann Harman, Talvin Momi, Harvey Wilkes). **Authors contribution statement:** RP conceived the study and designed the research approach. RP, KP, KW, RG, OA, LAH, TM and HW performed the experiments. RP, KP, KW, RP and OA analysed the data. RP and CDP wrote the first draft of the manuscript. All authors provided feedback and approved the final version of the manuscript.

**Keywords:** Agave, Wounds, Antibacterial, Biofilm

## Abstract

The global threat posed by antibiotic resistant infections highlights the urgent need to identify novel antimicrobials. In recent years, there has been a growing interest in revisiting and evaluating historical infection remedies for their antimicrobial activity. The *Agave* plant held significant social, religious, and medicinal value to the Mexica (commonly referred to as the Aztecs), as well as other Indigenous Mesoamerican communities. The use of Agave syrup mixed with salt was traditionally used in wound care as documented in the Florentine Codex, a sixteenth-century text authored by Spanish Franciscan friar Bernardino de Sahagún. The minimum inhibitory concentration was determined through the microbroth dilution method, whilst anti-biofilm activity was measured using crystal violet staining. In this current study, we demonstrated that commercially available Agave syrups primarily derived from the *Agave tequilana* species significantly inhibit the growth of Gram-positive and Gram-negative wound-associated pathogens in planktonic culture. The traditional practice of adding salt enhanced the efficacy against *Escherichia coli.* Furthermore, Agave syrups significantly inhibited biofilm formation across all six organisms tested, although their capacity to disrupt pre-formed biofilms appears to be species specific. Preliminary investigations into the underlying antibacterial mechanisms of Agave syrups suggest that it is likely due to a combination of factors, including their acidic pH, along with the presence of saponins, methylglyoxal, and the generation of hydrogen peroxide. This study contributes to the growing evidence that historical remedies like Agave syrup may be effective as new antimicrobial treatments.

## Introduction

Wound management is estimated to cost the NHS £8.3 billion per annum (1). The exposed subcutaneous tissue caused by trauma, surgery, burns, and ulcers favours bacterial colonisation (2, 3) with the most common species isolated from wounds being the Gram-positive skin commensal *Staphylococcus aureus,* and the Gram-negative organisms *Pseudomonas aeruginosa*, *Acinetobacter baumannii* and *Escherichia coli* (4). The ability of these organisms and others to form biofilms is known to delay wound healing and contribute to the development of chronic wound infections (5).

Biofilms are communities of bacterial aggregates which are embedded in an extracellular matrix rich in polysaccharides, proteins, and DNA (6, 7). This matrix provides protection against the host’s immune response and reduces the penetration of antibiotics (8). Thus, biofilm-associated infections are known to typically require antibiotic concentrations that are 100-1000 times higher than those required to treat the same microorganisms grown in free-floating planktonic culture (9, 10). Due to problems posed by biofilm infections and critical concerns around antibiotic resistance (11), there is an increasing need to identify novel antimicrobial agents.

Over recent decades, there has been an increasing interest in looking to the past to identify and scientifically evaluate pre-modern and traditional remedies for infection (12, 13). This has led to the emergence of successful interdisciplinary collaborations between microbiologists, data scientists and historians (14, 15). Bald’s eyesalve, an eleventh-century Old English remedy used traditionally for the treatment of “wens” or styes, has demonstrated powerful antimicrobial and anti-biofilm activity against *S. aureus, A. baumannii* and *Neisseria gonorrhoea* (11, 16, 17). The fields of ethnopharmacology and ethnobotany have also proved productive, working with Indigenous peoples and medicinal traditions to share traditional knowledge. An ethnohistoric study of sixteenth-century Mexican traditional medicine identified more than 3,000 plants possessing bioactive compounds that exhibit a diverse range of activities, which include antimicrobial, anti-inflammatory and insecticidal effects (Béjar *et al.,* 2000).

With respect to Mesoamerica specifically, extracts from Maya medicinal plants have been shown to exhibit bactericidal and anti-biofilm activity against *S. aureus,* including drug-resistant strains (18). Moreover, bark extracts from the Maya medicinal plant *Matayba oppositifolia* exhibited antibacterial activity against both carbapenem-resistant *Klebsiella pneumoniae* and *A. baumannii* (19). Among the Aztec-Mexica, *Artemisia mexicana* subsp. *ludoviciana*, a type of wormwood known in Nahuatl as *iztauhyatl*, was traditionally used for the treatment of digestive complaints and is still widely used for treating gastrointestinal conditions, pain and diabetes (20). Recent studies have confirmed its efficacy against not only diabetes (21) but also various microorganisms responsible for gastrointestinal issues, including *Helicobacter pylori* (22).

The *Agave* plant with its central pina surrounded by large spiral leaves (23) is known to have played a significant role in Aztec-Mexica society, from the production of alcoholic pulque, textiles, paper and roofing materials, to its use for medicinal purposes (24, 25). Commonly known in Spanish as *maguey* – likely derived from the Taíno *mawei* - this family of succulents was called *metl* by the Nahua (Nahuatl-speaking) peoples of Central Mexico (26, pp. 122-3). It is not always possible to distinguish specific genera in ethnographic sources, but one of the earliest Aztec-Mexica references to the use of *Agave* in wound management is contained within the Florentine Codex, a sixteenth-century text composed in collaboration with Indigenous informants and partners by the Spanish friar Bernardino de Sahagún. According to the friar’s Indigenous informants, wounds were treated with juice extracted from ‘well cooked’ *tlacametl* (*Agave atrovirens*) mixed with salt. Here, syrup was extracted from the *Agave* plant after being boiled or cooked over a fire (24, 27, Book 11, pp. 179). Head injuries were seen as particularly suitable for treatment with maguey sap; a mere wound was washed in urine, followed by ‘maguey leaf sap’, while a broken skull was joined with a bone awl, before being covered either with maguey sap or a freshly grated green maguey leaf (28, Book 10, pp. 140-141).

Davidson and Ortiz de Montellano (1983) were the first to empirically test a commercial Agave sap “Heart O Maguey” obtained from the leaves of *Agave atrovirens* (*A. atrovirens*). Here, they demonstrated through traditional disc diffusion assays that the sap, with and without the traditional addition of salt inhibited the growth of several clinically relevant pathogens, including *S. aureus, P. aeruginosa* and *E. coli.* Moreover, high-fructose Agave syrups have been shown to exhibit antimicrobial activity against both *E. coli* DH5 and *Bacillus subtilis* 168 (29). Methanolic leaf extracts of *Agave americana* (*A. americana*) were shown to possess antimicrobial properties against *S. aureus*, *P. aeruginosa*, and *Salmonella typhi* strains through disc diffusion experiments (30). More recently, Agave fructans, a prebiotic from *Agave tequilana* (*A. tequilana*), was shown to inhibit the growth and biofilm formation of a range of multi-drug-resistant isolates of *S. aureus* involved in bovine mastitis (31). However, further studies into the ability of Agave syrups to inhibit wound-associated pathogens are lacking, as are investigations into its minimum inhibitory concentration and its anti-biofilm activity (12).

This study aimed to build upon previous findings to evaluate the antibacterial and anti-biofilm activity of Agave syrup against a range of wound-associated pathogens known to cause infections in humans. Initial experiments sought to determine the minimum inhibitory concentrations (MIC) of Agave syrups, and determine biofilm inhibition and eradication activity, as well as conduct preliminary studies to investigate its potential mechanisms of action. It is important to clarify that our work did not seek to replicate the exact methods employed by the Aztec-Mexica, such as the use of freshly crushed leaves from the *A. atrovirens, Agave mexicana* (*A. mexicana*) and *Agave potatorum* (*A. potatorum*) species (24). Instead, we investigated the antimicrobial activity of five commercially available syrups, principally derived from *A. tequilana* (commonly known as blue agave), with and without the traditional addition of salt. This allowed us to honour the historical use of Agave syrup by Indigenous communities for the treatment of wounds, whilst utilising a standardised, reproducible, and globally available test material.

## Materials and Methods

### Bacterial strains and growth conditions

All bacterial strains used in this study are listed in Table 1. Strains were stored at −80 °C in 50% (v/v) glycerol/water. All cultures were streaked from frozen stocks onto Luria Bertani (LB) agar plates (Fisher Scientific, UK) and grown aerobically at 37 °C for 48 h. To ensure a pure culture, strains were Gram stained and selective agar plates were utilised.

**Table 1.** Bacterial strains used in this study.

|  | Species | Strain |
| --- | --- | --- |
| Gram-positive | *Staphylococcus aureus* | ATCC 6538 |
|  | *Staphylococcus epidermidis* | NCTC 13360 |
|  | *Streptococcus pyogenes* | NCTC 126916 |
| Gram-negative | *Pseudomonas aeruginosa* | PAO1 |
|  | *Escherichia coli* | NCTC 12241 |
|  | *Acinetobacter baumannii* | NCTC 12156 |

### Media

Luria Bertani (LB) agar, LB broth and Mueller Hinton Broth (MHB) were obtained from Fisher Scientific, UK. Artificial wound media consisted of equal volumes of sterile peptone water (Fisher Scientific, UK) and heat-inactivated foetal bovine serum (Gibco, UK) (11, 32).

### Agave syrups

A total of five commercially available Agave syrups were included in this study; details of each are shown in Table 2. Agave syrups 1-3 and 5 were derived from the *A. tequilana* species, whilst Agave 4 was derived from a mixture of *A. tequilana* and *Agave salmiana.* Manufacturers were contacted to ascertain the part of the Agave plant used in the production of each syrup. Agave 4 is produced from the leaves, whilst Agave 5 is from the central pina. The manufacturers of Agave syrups 1-3 did not respond to inquiries and consequently the parts of the *Agave* plant used to produce syrups 1-3 remain unidentified.

**Table 2.** Agave syrups and Manuka honey used in this study.

| Treatment | Name | Agave species | Source | Part of the plant used |
| --- | --- | --- | --- | --- |
| Agave 1 | Organic Agave plant syrup | *Agave tequilana* | Tate and Lyle | Unknown |
| Agave 2 | Sirope De Agave | *Agave tequilana* | Naturitas | Unknown |
| Agave 3 | Organic dark blue Agave syrup | *Agave tequilana* | Kirkland Signature | Unknown |
| Agave 4 | Organic Agave Nectar | *Agave tequilana* & *Agave salmiana* | Vitanics | Leaves |
| Agave 5 | Organic raw Agave syrup | *Agave tequilana* | Indigo Herbs | Pina |
| Manuka Honey | MGO70 Manuka Honey | New Zealand Manuka Honey | Manuka Doctor, UK | N/A |

Each Agave syrup was stored in the dark at room temperature. Prior to testing, 90% (w/v) stock solutions of each syrup were prepared by weighing 18 g of Agave syrup in a sterile 50 mL tube and bringing the volume up to 20 mL using sterile deionised water. A 90% (w/v) syrup was chosen to make the syrup easier to pipette, but it is also not known how concentrated the syrup was when used for traditional medicinal purposes. A 90% (w/v) stock solution of Manuka honey (Manuka Doctor, UK) was used as a positive control, due to its well documented antimicrobial activity (33, 34). These 90% (w/v) stock solutions were stored at 4°C. Stock solutions of Agave syrup with salt and Manuka honey with salt were prepared as described above, with the addition of 2 g of sodium chloride (Fisher Scientific, UK).

All 90% (w/v) solutions of Agave syrup and Manuka honey were sterilised by exposure to UV radiation for 30 minutes (ThermoFisher Cabinet, fitted with a 2537 7 Å, 8-W UV bulb). To confirm sterility, a sterile cotton swab was dipped into each solution and streaked across the surface of an LB agar plate. Plates were incubated for 24 hrs at 37 °C prior to being visually inspected to confirm sterility indicated by the absence of visual bacterial growth.

### Determination of minimum inhibitory concentration (MIC) and minimum bactericidal concentration (MBC)

The MIC of the five Agave syrups and Manuka honey (positive control) was determined as described by Furner-Pardoe *et al.* (2020) in double strength MHB. Sterile deionised water was used as a negative control. Following overnight growth on LB agar, three to five morphologically similar colonies of each organism were added to sterile 1x phosphate-buffered saline (PBS) and adjusted to a 0.5 McFarland solution (OD_600_ 0.08-0.1) using a UV/Visible spectrophotometer 6315 (Beckman Instrument Ltd, UK). The bacterial suspension was then diluted 1:100 in fresh 2x MHB, prior to 100 µL of the bacterial suspension being added to respective wells of a sterile round bottom 96-well plate containing 100 µL of the diluted Agave syrup or Manuka honey. Each stock concentration of 90% (w/v) Agave was two-fold diluted when added to the bacterial inoculum, resulting in the final concentrations: 45%, 22.5%,11.25%, 5.63%, 2.81% and 1.41% (w/v) respectively.

Plates were incubated for 18 h at 37 °C prior to being visually inspected. The lowest concentration of treatment which prevented visible growth of each organism was recorded as the MIC. To determine if the antimicrobial activity of Agave was bacteriostatic or bactericidal, 25 µL from each well where visible growth was inhibited were added to the surface of LB agar plates and incubated at 37 °C for 24 h. Agar plates with visible colony growth following incubation were recorded as having bacteriostatic activity, whilst no growth was recorded as bactericidal activity (MBC). All experiments were repeated using Agave syrup and Manuka honey stock concentrations mixed with 2 g of salt.

### Biofilm Inhibition Assay

The ability of Agave syrup to inhibit bacterial biofilm production was determined using the crystal violet staining technique, adapted from (35). Two-three colonies of a respective organism were incubated in artificial wound media on an orbital shaker at 37 °C for 4 hrs. The cultures were then diluted to OD_600_ 0.08-0.1 in fresh artificial wound media. A total of 100 µL of each culture were added to the wells of a sterile flat bottomed 96-well plate, along with 100 µL of 90% (w/v) Agave or Manuka honey. The 96-well plates were incubated aerobically at 37°C for 24 h under static conditions. Following this, well contents were aspirated and discarded, and biofilms were gently washed using 200 µL of 1x PBS before being heat fixed at 60 °C for 30 minutes.

Plates were then stained with 200 µL of 1% (w/v) crystal violet for 10 minutes and washed twice in deionised water. After drying overnight, 200 µL of 30% (v/v) acetic acid was added to each well for 10 minutes to solubilise the stain, after which it was transferred to a new 96-well plate and read at OD_550_ on a plate reader (ThermoScientific – Multiskan Sky). Experiments were repeated using Agave syrup and Manuka honey mixed with 2g of salt. Biofilm inhibition was reported as a percentage of the untreated control to account for variability in baseline biofilm formation and to facilitate comparison across independent experiments.

### Disruption of pre-formed biofilms

The ability of Agave syrup to eradicate pre-formed biofilms was determined using the crystal violet staining technique, adapted from (35). Two-three colonies of a given organism were incubated in artificial wound media on an orbital shaker for 4 h, at 37 °C. The incubated cultures were then diluted to OD_600_ 0.08-0.1 in fresh artificial wound media, with a total of 100 µL of each culture then being added to the wells of a sterile flat bottomed 96-well plate. Plates were incubated aerobically at 37°C for 24 h under static conditions. Following this, 100 µL of 90% (w/v) Agave or Manuka honey was added to the wells of the 96-well plate, prior to the plate being incubated for a further 24 h. Biofilms were then gently washed using 200 µL of 1x PBS and heat fixed at 60 °C for 30 minutes (VWR Digital Mini Incubator). Plates were then stained with 200 µL of 1% (w/v) crystal violet for 10 minutes and washed twice in deionised water. After drying overnight, 200 µL of 30% (v/v) acetic acid was added to each well for 10 minutes to solubilise the stain, prior to the stain being transferred to a new 96-well plate. The plate was then read at OD_550_ (ThermoScientific – Multiskan Sky). Biofilm inhibition was reported as a percentage of the untreated control to account for variability in baseline biofilm formation and to facilitate comparison across independent experiments.

### Screening for the presence of saponins

The presence of saponins within each Agave syrup and Manuka honey was determined following the method as described by Shegute and Wasihun (2020). A total of 1g of Agave syrup (with or without the addition of 2g salt) was added to 9 mL of sterile deionised water within a 20 mL universal tube. The tube was then shaken vigorously for 15 seconds, prior to being allowed to stand in a rack for 20 minutes. The presence and persistence of a top layer foam upon warming was confirmed as a positive result for the presence of saponins, whilst the absence of foam was a negative result.

### Measuring of the pH of the Agave

The pH of each 90% (w/v) Agave syrup and Manuka honey was measured using a pH glass electrode attached to a Hanna H18424 pH meter. The probe was calibrated prior to use, using three-point calibration solutions (pH 4, pH 7 and pH 10).

### Determining hydrogen peroxide production

The presence of hydrogen peroxide within each of the 90% (w/v) Agave syrups and Manuka honey was determined using an Amplex Red hydrogen peroxide kit (Fisher Scientific, UK) following the manufacturer’s instructions. A 20 µM hydrogen peroxide stock solution was serially diluted two-fold to construct a standard curve. 50 µL of the standard curve samples, controls, Agave syrups and Manuka honey were added to wells of a microtiter plate. Plates were incubated for 30 minutes prior to the absorbance being read at a wavelength of 560 nm on a plate reader (ThermoScientific – Multiskan Sky). Absorbance values were subtracted from wells containing Agave syrup and 1x reagent buffer.

### Methylglyoxal determination

The concentration of methylglyoxal within each Agave syrup and the Manuka honey control was determined using the *N*-acetyl-L-cysteine (NAC) assay as described previously by (36). Methylglyoxal standards were produced using a total volume of 1 mL, consisting of 20 µL of 500 mM *N*-acetyl-L-cysteine, 100 mM methylglyoxal (5-50 µL) and 100 mM NaH_2_PO_4_ (pH 7.0). To determine the methylglyoxal concentration within each Agave syrup, 50 µL of each Agave syrup was mixed with 930 µL of 100 mM NaH_2_PO_4_ (pH 7.0) and 20 µL of 500 mM *N*-acetyl-L-cysteine. Samples were incubated for 10 minutes at room temperature and regularly inverted, prior to being read at 288 nm on UV/Visible spectrophotometer 6315 (Beckman Instrument Ltd, UK). To correct for background absorbance, absorbance readings of Agave and Manuka honey alone were subtracted from the absorbance readings obtained for methylglyoxal quantification.

### Statistical analysis

All results unless otherwise stated are expressed as mean ± S.E.M. The number of independent and technical repeats is detailed within each figure legend. All statistical analyses were performed using GraphPad Prism 10.2 software (GraphPad, La Jolla, CA, USA) with significance being set to *p*<0.05. Any outliers were identified and removed using the ‘Identify Outliers’ function on GraphPad. The specific statistical tests and *post-hoc* tests used for each experiment are described in the figure legends.

## Results

The antimicrobial activity of the five Agave syrups and Manuka honey positive control was tested against six wound-associated pathogens, as shown in Table 3.

**Table 3.** The minimum inhibitory concentration (MIC) of five Agave syrups and one Manuka honey with and without salt against three Gram-positive and three Gram-negative organisms following growth in double strength MHB. Results shown are the modal MIC from three independent experiments (*N*=3), each performed in duplicate.

| Treatment | *S. aureus*<br>MIC (% w/v) | *S. epidermidis*<br>MIC (% w/v) | *S. pyogenes*<br>MIC (% w/v) | *A. baumannii*<br>MIC (% w/v) | *E. coli*<br>MIC (% w/v) | *P. aeruginosa*<br>MIC (% w/v) |
| --- | --- | --- | --- | --- | --- | --- |
| Agave 1 | 22.5 | 45 | 45 | 11.25 | 45 | 22.5 |
| Agave 1 with salt | 45 | 45 | 45 | 11.25 | 45 | 22.5 |
| Agave 2 | 22.5 | 45 | 45 | 11.25 | 45 | 22.5 |
| Agave 2 with salt | 45 | 45 | 45 | 11.25 | 22.5 | 22.5 |
| Agave 3 | 45 | 45 | 45 | 11.25 | 45 | 22.5 |
| Agave 3 with salt | 45 | 45 | 45 | 11.25 | 45 | 22.5 |
| Agave 4 | 22.5 | 45 | 45 | 11.25 | 45 | 22.5 |
| Agave 4 with salt | 22.5 | 45 | 45 | 11.25 | 22.5 | 22.5 |
| Agave 5 | 22.5 | 45 | 45 | 11.25 | 45 | 22.5 |
| Agave 5 with salt | 22.5 | 45 | 45 | 11.25 | 22.5 | 22.5 |
| Manuka Honey | 11.25 | 11.25 | 22.5 | 5.62 | 22.5 | 22.5 |
| Manuka Honey with salt | 11.25 | 11.25 | >45 | 5.62 | 22.5 | 22.5 |

The efficacy of Agave syrup against the Gram-positive organism *S. aureus* is shown in Table 3. The MIC for Agave 1 and 2 were 22.5% (w/v), with this increasing to 45% (w/v) following the addition of salt. The MIC for Agave 3 was 45% (w/v), whilst Agave 4 and 5 had an MIC of 22.5% (w/v), all of which were unaffected by the addition of salt. The positive control of Manuka honey exhibited the lowest MIC at 11.25% (w/v) with the MIC not being influenced by the addition of salt. All five Agave syrups exhibited an MIC of 45% (w/v) against *S. epidermidis* and this was not influenced by the addition of salt. As seen with *S. aureus*, Manuka honey had the lowest MIC of all the treatments tested at 11.25% (w/v) and the MIC was unchanged following the addition of salt. All five Agaves exhibited an MIC of 45% (w/v) when tested against *S. pyogenes*, with Manuka honey having the lowest MIC at 22.5% (w/v). The MICs for each of the Agaves remained unchanged following the addition of salt, however, this increased the MIC for Manuka honey to >45% (w/v).

Next, the efficacy of Agave syrup against the Gram-negative pathogen *A. baumannii* was determined. The MIC for all five Agave syrups was 11.25% (w/v) and this was unaffected following the addition of salt. Manuka honey exhibited the lowest MIC, with this being 5.62% (w/v) and this was unaffected by the addition of salt. With respect to *E. coli*, the MICs of all five Agave syrups was 45% (w/v) with Manuka honey having the lowest MIC of 22.5% (w/v). The addition of salt did not affect the MIC of Agave 1 and 3 or Manuka honey, whilst it reduced the MIC for Agaves 2, 4 and 5. Finally, with respect to *P. aeruginosa*, the MIC for each of the five Agave syrups and Manuka honey was 22.5% (w/v) and this was not influenced by the addition of salt.

Small volumes of culture from each of the MIC wells were pipetted onto LB agar and incubated overnight at 37 °C to determine bacterial viability. Results from these experiments indicate that Agave syrup is bacteriostatic in its activity, as growth was seen following inoculation onto LB agar.

### Inhibition of biofilm formation

As mentioned previously, chronic wound infections are typically characterised by the biofilm mode of growth. To address this, we sought to determine whether Agave syrup could inhibit biofilm production by Gram-positive wound associated pathogens grown in artificial wound media. As shown in Figure 2A, Agave 1, Agave 4, Agave 5 and Manuka honey significantly inhibited *S. aureus* biofilm formation compared to the untreated control (*p*<0.001, *p*=0.0231, *p*=0.0034 and *p*=0.0047 respectively). The addition of salt did not significantly enhance this anti-biofilm effect further compared to the Agave syrup alone. In contrast, Agave syrups 2 and 3 alone did not significantly reduce *S. aureus* biofilm formation compared to the untreated control, however, when mixed with salt, this led to a significant reduction in biofilm biomass (*p*<0.0152 and *p*<0.001 respectively).

**Figure 1.** Investigating the antibacterial and anti-biofilm activity of commercial Agave syrups with and without salt, based on their historic and Indigenous use in wound care. Initial experiments sought to determine the minimum inhibitory concentration (MIC) of five Agave syrups against three Gram-positive and three Gram-negative wound associated pathogens following growth in planktonic culture. Bacterial viability was determined through plating onto agar. Anti-biofilm activity was determined through crystal violet staining. The physicochemical properties of each syrup were also determined, screening for saponins, pH, hydrogen peroxide (H_2_O_2_) and methylglyoxal (MGO). Figure for the graphical abstract was created using Biorender (https://BioRender.com).

**Figure 2.** Agave syrups and Manuka honey, with and without salt, significantly reduce *S. aureus*, *S. epidermidis* and *S. pyogenes* biofilm formation. Biofilm formation is expressed as a percentage of the untreated control. Results show the mean biofilm formation ± S.E.M for three independent experiments (*N*=3), each performed in triplicate. Statistical differences were determined using a two-way ANOVA with Tukey’s *post-hoc*. Asterisks indicate significance vs the control, whilst lines with asterisks indicate significance between a given Agave with and without salt. \**p*<0.05, \*\**p*<0.01 and \*\*\**p*<0.001. A1-A5 = Agave 1-5. + NaCl = with salt. H = Manuka Honey.

For *S. epidermidis* biofilm formation (Figure 2B), all five Agave syrups and Manuka honey significantly reduced biofilm formation compared to the untreated control (Agave 1, 3 and 5: *p*<0.001; Agave 2: *p*=0.0039; Agave 4: *p*=0.0085 and Manuka honey: *p*= 0.0010). The addition of salt did not enhance this effect, except for Agave 4, which showed a further significant reduction (*p*=0.0089).

Finally, each Agave syrup significantly inhibited *S. pyogenes* biofilm formation (Figure 2C) compared to the untreated control (Agave 1-4: *p*<0.001; Agave 5: *p*=0.0023). Interestingly, Manuka honey did not significantly inhibit *S. pyogenes* biofilm formation compared to the untreated control, whilst the addition of salt to Manuka honey resulted in a significant reduction in the biofilm biomass compared to the untreated control (*p*=0.0252).

Next, we investigated whether Agave syrup could inhibit biofilm formation by Gram-negative, wound-associated pathogens grown in artificial wound media. As shown in Figure 3A, all Agave syrups and Manuka honey significantly reduced biofilm biomass of *A. baumannii* (*p*<0.001) compared to the untreated control. Similarly, each Agave syrup and Manuka honey significantly inhibited *E. coli* biofilm formation (Figure 3B) (*p*<0.001) and *P. aeruginosa* biofilm formation (Figure 3C) compared to the untreated control (*p*<0.001). The addition of salt had no further impact on this inhibitory effect, except for Agave 1 with salt and Agave 3 with salt, which demonstrated enhanced activity against *E. coli* (*p*=0.0312 and *p*=0.0445).

**Figure 3.** Agave syrups and Manuka honey, with and without salt, significantly reduce *A. baumannii, E. coli* and *P. aeruginosa* biofilm formation. Biofilm formation is expressed as a percentage of the untreated control. Results show the mean biofilm formation ± S.E.M for three independent experiments (*N*=3), each performed in triplicate. Statistical differences were determined using a two-way ANOVA with Tukey’s *post-hoc*. Asterisks indicate significance vs the control, whilst lines with asterisks indicate significance between a given Agave with and without salt. \**p*<0.05, \*\**p*<0.01 and \*\*\**p*<0.001. A1-A5 = Agave 1-5. + NaCl = with salt. H = Manuka Honey.

### Eradication of pre-formed biofilms

After establishing that Agave syrups could inhibit biofilm production, we then evaluated their ability to eradicate pre-formed biofilms produced by Gram-positive pathogens. All Agave syrups and Manuka honey both with and without salt failed to significantly reduce or eliminate established biofilms produced by *S. aureus* (Figure 4A)*, S. epidermidis* (Figure 4B) and *S. pyogenes* (Figure 4C). For both *S. aureus* and *S. epidermidis*, the addition of Agave was associated with a non-significant trend toward increased biofilm formation, particularly following the addition of salt.

**Figure 4.** Mature biofilms of *S. aureus, S. epidermidis* and *S. pyogenes* are not affected following treatment with Agave syrups and Manuka honey. Results show the mean biofilm formation ± S.E.M for three independent experiments (*N*=3), each performed in triplicate. Statistical differences were determined using a two-way ANOVA with Tukey’s *post-hoc*. Asterisks indicate significance vs the control, whilst lines with asterisks indicate significance between a given Agave with and without salt. A1-A5 = Agave 1-5. + NaCl = with salt. H = Manuka Honey.

The efficacy of Agave syrup and Manuka honey to eradicate pre-formed biofilm produced by Gram-negative pathogens was also determined. All Agave syrups caused significant reductions in biofilm biomass produced by *A. baumannii* (Figure 5A) compared to the untreated control (*p*<0.001), and the addition of salt did not enhance this further. Whilst Manuka honey reduced the biofilm biomass of *A. baumannii* compared to the untreated control (*p*=0.0013), the addition of salt did not. Similarly, all five Agave syrups, both with and without salt significantly reduced the bacterial biomass of pre-formed *E. coli* biofilms (Figure 5B) compared to the untreated control (*p*<0.001). Manuka honey significantly reduced the biofilm biomass of *E. coli* compared to the untreated control as did Manuka honey with salt (*p*<0.001). However, neither Agave syrups nor Manuka honey had any significant effect on pre-formed *P. aeruginosa* biofilms, with the addition of salt failing to elicit any further effect.

**Figure 5.** Agave syrups and Manuka honey, both with and without salt, significantly reduce pre-formed biofilms of Gram-negative species in an isolate-dependent manner. Biofilm formation is expressed as a percentage of the untreated control. Results show the mean biofilm formation ± S.E.M for three independent experiments (*N*=3), each performed in triplicate. Statistical differences were determined using a two-way ANOVA with Tukey’s *post-hoc*. Asterisks indicate significance vs the control, whilst lines with asterisks indicate significance between a given Agave with and without salt.. \**p*<0.05, \*\**p*<0.01 and \*\*\**p*<0.001. A1-A5 = Agave 1-5. + NaCl = with salt. H = Manuka Honey.

To investigate the potential mechanisms underlying the antimicrobial activity of each Agave syrup, a series of experiments was conducted. These included screening for saponins, measuring the pH, assaying hydrogen peroxide production, and determining methylglyoxal concentration.

### Screening for the presence of saponins

Saponins were detected in all five Agave syrups and Manuka honey, as indicated by the persistence of foam over time (Figure 6A). However, following the addition of salt, saponins were only visually detected in Manuka honey (Figure 6B).

**Figure 6.** Screening for the presence of saponins within Agave syrup and Manuka honey with and without the addition of salt. Figure 6A shows the screening of saponins in Agave syrups (A1-A5) and Manuka honey (H). Figure 6B shows the screening of the Agave syrups and Manuka honey containing salt. Images are representative of three independent experiments (*N*=3).

### Determining the pH of the Agave syrups

Next, we sought to determine the pH of the Agave syrups and Manuka honey. As shown in Table 4, each of the Agave syrups and Manuka honey were acidic, with a low mean pH, ranging from 3.28-4.38.

**Table 4.** The mean pH of Agave syrup, with and without the addition of salt. Experiments were performed twice (*N*=2), with each measurement being taken in duplicate.

| Treatment | Mean pH | Range |
| --- | --- | --- |
| Agave 1 | 4.38 | 4.31-4.45 |
| Agave 2 | 4.22 | 4.08-4.35 |
| Agave 3 | 3.97 | 3.91-4.02 |
| Agave 4 | 3.63 | 3.62-3.64 |
| Agave 5 | 3.28 | 3.15-3.41 |
| Manuka Honey | 3.47 | 3.22-3.72 |

### Measuring hydrogen peroxide production

Given that a known antimicrobial mechanism of Manuka honey is the generation of hydrogen peroxide, we sought to determine the concentration of hydrogen peroxide (H_2_O_2_) within each of the Agave syrups and Manuka honey. As shown in Figure 7, whilst H_2_O_2_ was detected in each of the five Agave syrups tested (mean ranging between 5.19-8.82 μM), the concentration detected within Manuka honey was significantly higher (mean 35 μM) (*p*<0.001). There were no significant differences in the hydrogen peroxide concentration across the five Agave syrups.

**Figure 7.** The concentration of hydrogen peroxide (H_2_O_2_) detected within each of the Agave syrups, compared to the honey control. Hydrogen peroxide production was assayed using an Amplex Red ® kit in accordance with the protocol provided. A1 = Agave 1, A2 = Agave 2, A3 = Agave 3, A4 = Agave 4, A5 = Agave 5 and H = Manuka honey. Results show the mean concentration of H_2_O_2_ ± S.E.M for three independent experiments (*N*=3), each performed in triplicate. Statistical differences were determined using a one-way ANOVA with Tukey’s *post-hoc.* \*\*\**p*<0.001.

### Methylglyoxal Determination

Finally, the concentration of methylglyoxal (MGO) within each of the Agave syrups and Manuka honey was determined using the *N*-acetyl-L-cysteine (NAC) assay. As shown in Figure 8, the Agave syrups varied in their MGO concentration, with the concentration in Agave 5 being below the limit of detection. The concentration of MGO within Manuka honey was significantly higher compared to each of the Agave syrups (*p*<0.001). Agave 2 had a significant higher MGO concentration compared to the other four Agave syrups (*p*=0.055 vs. Agave 1; *p*=0.0029 vs. Agave 3; *p*<0.001 vs. Agave 4 and *p*<0.001 vs. Agave 5).

**Figure 8.** The concentration of methylglyoxal (MGO) detected within each treatment. The concentration of MGO within each of the treatments was determined using the *N*-acetyl-L-cysteine (NAC) assay. A1 = Agave 1, A2 = Agave 2, A3 = Agave 3, A4 = Agave 4, A5 = Agave 5 and H = Manuka honey. Results show the mean concentration of MGO ± S.E.M for three independent experiments (*N*=3). Statistical differences were determined using a one-way ANOVA with Tukey’s *post-hoc. **p*<0.01, \*\*\**p*<0.001.

## Discussion

This study aimed to assess the antibacterial and anti-biofilm activity of Agave syrup as a traditional remedy for wound management. The antibacterial activity of Agave syrup was first reported by Davidson and Ortiz de Montellano (1983), through traditional disc diffusion experiments. Here, they demonstrated that commercially available maguey sap from *Agave atrovirens* produced a zone of inhibition, inhibiting the growth of six clinically significant pathogens, including *S. aureus*, *P. aeruginosa* and *E. coli.* Since then, several studies have explored the antimicrobial activity of Agave against a range of bacterial and fungal pathogens using methanol and ethanol extracts and rotatory evaporation to concentrate the supernatants (30, 37, 38).

In this study, all five commercially available Agave syrups exhibited broad bacteriostatic activity against both Gram-positive and Gram-negative pathogens following their growth in planktonic culture (as shown in Table 3). Two of the Gram-negative organisms, *P. aeruginosa* and *A. baumannii*, exhibited the greatest sensitivity to Agave syrup, evidenced by their low MIC values. This finding aligns with Davidson and Ortiz de Montellano, (1983) who demonstrated that the four Gram-negative pathogens included in their study -*P. aeruginosa, E. coli, Salmonella paratyphi* and *Shigella sonnei* - exhibited the largest zones of inhibition compared to the two Gram-positive cocci *S. aureus* and *Sarcina lutea*. The enhanced susceptibility of Gram-negative bacteria to the antibacterial effects of Agave syrup may be explained, in part, to having a thinner peptidoglycan cell wall, which might otherwise provide a degree of protection against the syrup’s antimicrobial activity (39).

The traditional addition of salt did not enhance the antibacterial efficacy of Agave syrup for five of the six pathogens tested following growth in planktonic culture. The exception was *E. coli*, where an increased MIC was observed when exposed to Agave with salt (Table 3). This effect may be explained in part due to *E. coli’*s low tolerance to salt (40), however further investigation is needed to confirm this. These findings are largely in agreement with Davidson and Ortiz de Montellano, (1983), who demonstrated that salt did not enhance the zone of inhibition for any of the organisms tested, except for *S. aureus.* The lack of enhancement in the MIC for *S. aureus* in this study may be due to the organism’s well-documented tolerance to high salt concentrations (41–43) and differences in the specific strain used, as well as the species of Agave from which the syrup was extracted.

In addition to testing the antimicrobial activity of Agave syrup against organisms grown in free-swimming planktonic culture, it is important to determine their efficacy against biofilms. Biofilms are composed of exopolysaccharide, DNA and proteins and are associated with chronic wound infections (44, 45). Whilst the anti-biofilm activity of Agave fructans has been reported in the treatment of bovine mastitis (31) experiments investigating the anti-biofilm activity of Agave syrups against human pathogens are limited (12).

High concentrations of Agave syrup (90% w/v) were used for the biofilm experiments as organisms growing in a biofilm are known to exhibit an increased tolerance to antibiotics at concentrations greater than 1,000 times that which is needed to treat planktonic cultures (10). The findings within this study demonstrate that Agave syrup significantly inhibits biofilm formation of all six wound-associated pathogens compared to the untreated control (Figures 2 and 3). This effect is likely due in part to Agave’s ability to inhibit bacterial growth, as observed in the MIC experiments (Table 3). The addition of salt to Agave 4 further enhanced its inhibitory effect upon *S. epidermidis* biofilm formation (Figure 2), whilst the addition of salt to Agave 1 and Agave 3 enhanced its inhibitory effect against *E. coli* biofilm formation (Figure 3). No other enhancements were seen when salt was added. Whilst these findings warrant further study, the ability of Agave syrup inhibiting biofilm formation is a key finding, as biofilm formation plays an essential role in the persistence and severity of wound infections (46, 47). As shown in Figures 4 and 5, the ability of Agave to eradicate already established biofilms is poor, particularly against the three Gram-positive organisms. The addition of Agave appeared to promote an increase in biofilm biomass for both *S. aureus* and *S. epidermidis,* although this was not significant. Increasing salt concentrations have been shown to increase biofilm formation of *S. aureus* (48, 49), along with *S. epidermidis* (50). Only *A. baumannii and E. coli* exhibited significant reductions in biofilm, whilst pre-formed biofilm by *P. aeruginosa* was unaffected by the addition of Agave syrup or Manuka honey. We hypothesise that this could be due to the mechanistic differences in biofilm formation by Gram-negative organisms (51, 52). In terms of the Agave syrup, it is likely that the established biofilm itself is providing protection against the antimicrobial mechanisms. Thus, the efficacy of Agave syrup in treating established biofilms within chronic wound infections is likely to be reduced.

Manuka honey was used in this study as a positive control, as its antimicrobial mechanisms have been widely studied and are believed to be due to a combination of different factors (53, 54). This includes its low pH, the presence of methylglyoxal, the production of hydrogen peroxide, its high osmolarity and the presence of the antimicrobial peptide, bee defensin-1 (55–57). To better understand the antimicrobial mechanisms of Agave syrup, the presence of saponins, its pH, the production of hydrogen peroxide and the presence of methylglyoxal within each of the Agave syrups was determined.

We propose that the broad bacteriostatic activity of the Agave syrups may be attributed in part to Agave species producing saponins (58, 59), which play a role in protecting plants against pathogens (60). We showed through a simple assay that each of the Agaves exhibited saponin activity through the formation of stable foam (Figure 6), where the detergent-like properties of saponins are believed to damage bacterial membranes (61). The size of the foam layer was vastly reduced in those Agaves where salt was added, which is likely due to an increase in the surface tension of water, reducing the stability of the foam (62).

It is also likely that the acidic pH of the Agave syrups which ranged from 3.28-4.38 also contributes to its antimicrobial activity (Table 4). Organisms such as *S. aureus* require a minimum pH of 4.0 for growth (63), whilst *E. coli, P. aeruginosa* and *S. epidermidis* require a minimum pH of 4.3, 4.4 and 4.5 respectively (56). The growth of *S. epidermidis* has previously been shown to be reduced at a pH of 4.0 (64).

It has been previously reported that Agave syrups produce hydrogen peroxide following their dilution in water (29). Hydrogen peroxide is a well-documented antimicrobial compound within honey (65), where the production of reactive oxygen species is known to damage bacterial membranes and proteins (66–68). The detection of hydrogen peroxide within each of the Agave syrups (Figure 7) is supported by Escobosa *et al.,* (2014), and is at a concentration significantly lower than that of Manuka honey. Thus, hydrogen peroxide may be an important factor in the antimicrobial activity of Agave syrups.

Commercial Agave syrups have been shown to contain methylglyoxal (29) and the antimicrobial activity of Manuka honey is largely attributed to this dicarbonyl compound, which can disrupt both protein and DNA synthesis, as well as damage the membrane (55, 69–71). Four of the five Agave syrups tested had detectable concentrations of methylglyoxal, which were significantly lower than that of the Manuka honey (Figure 8). Methylglyoxal has also been shown to reduce the expression of fimbriae, reduce bacterial adhesion, as well as disrupt the structure of bacterial flagella, thus reducing bacterial motility (72).

It is also likely that the antimicrobial activity of Agave may be explained in part due its high sugar content which is evident on the ingredient label of each syrup (29, 73). Agave is known to be high in fructose (73, 74), with high concentrations of fructose in honey and sugar solutions having been shown to inhibit *P. aeruginosa* biofilm formation (75). Gram-negative bacteria have previously been shown to be more susceptible than Gram-positive organisms to the osmotic effects of honey (34, 53).

It is highly unlikely that the antimicrobial activity of Agave syrup is due to a single factor, but rather a combination of factors. Bacteria such as *P. aeruginosa* are known to be able to detoxify methylglyoxal (54, 76), whilst many bacteria produce catalase to breakdown hydrogen peroxide (68). Furthermore, it is known that the antimicrobial efficacy of the Old English remedy Bald’s eyesalve is dependent upon the entire mixture, as opposed to a single ingredient or compound (11).

### Limitations and future work

There are several limitations associated with this study that could be addressed with future work. Each of the commercial Agave syrups used were derived from the species *A. tequilana,* with only Agave 4 containing syrup extracted from both *A. tequilana* and *A. salmiana.* Despite this variation in species, a wide variety of *Agave* species has been shown to exhibit antimicrobial activity following solvent extraction, including *A. lecheguilla*, *A. picta*, *A. scabra* and *A. lophantha* (37), *A. sialana* (38) and *A. americana* (30). Future work could benefit from the use of other Agave species to test the wider antimicrobial activity of the genus.

Furthermore, future work could benefit from using raw Agave leaves. Of the five Agave syrups used in this study, only Agave 4 is known to be produced from the leaves of the plant. Future research should focus on producing the syrup using raw Agave leaves, heating it over a flame, and adding salt, following the traditional production methods (24). Moreover, we do not know how much salt was added to Agave syrup when it was traditionally prepared, or how concentrated the syrup was prior to its application to a wound (24). Therefore, changes to the amount of salt added could be varied in future research, possibly in consultation with Mexican practitioners of traditional medicine.

Furthermore, whilst the *S. aureus* and *P. aeruginosa* reference strains in this study were initially isolated from wounds (77, 78), future work could seek to test the efficacy of Agave syrups against various clinical wound isolates. As researchers hypothesise the difference in biofilm reduction of the Gram-negative organisms is due to biofilm mechanisms (52, 79), future work should also seek to determine the mechanism behind the ability of Agave syrup to inhibit bacterial biofilm formation through analysing the expression of genes involved in biofilm formation. Finally, many chronic wound infections are known to be polymicrobial in nature (4) and previous reports using an *in vivo* wound model demonstrated that *P. aeruginosa* grown in a single species biofilm exhibited an increased sensitivity to gentamicin compared to its growth in mixed culture (80). Thus, future work would seek to determine the efficacy of Agave syrup in treating polymicrobial biofilms.

## Conclusion

This is the first time within the authors’ knowledge that the effect of Agave syrup upon human wound-pathogens and their biofilms has been documented. Despite the limitations outlined above, each of the commercially available Agave syrups primarily derived from *A. tequilana* demonstrated antimicrobial abilities against the six pathogens. Apart from *E. coli,* no changes were seen in the antimicrobial efficacy of Agave following the traditional addition of salt following growth in planktonic culture. In terms of biofilms, we note that Agave syrup can inhibit biofilm formation across the six organisms tested but the Agave syrup failed to disrupt pre-formed biofilms for each of the three Gram-positive organisms, in addition to the Gram-negative organism *P. aeruginosa.* The antimicrobial mechanisms of the Agave syrups appear to be due to a combination of factors, including their acidic pH, along with the presence of saponins, methylglyoxal, and hydrogen peroxide. This study demonstrates that Indigenous knowledge and methods, like the use of Agave syrup in medicine, may offer effective new and alternative antimicrobial treatments.

---

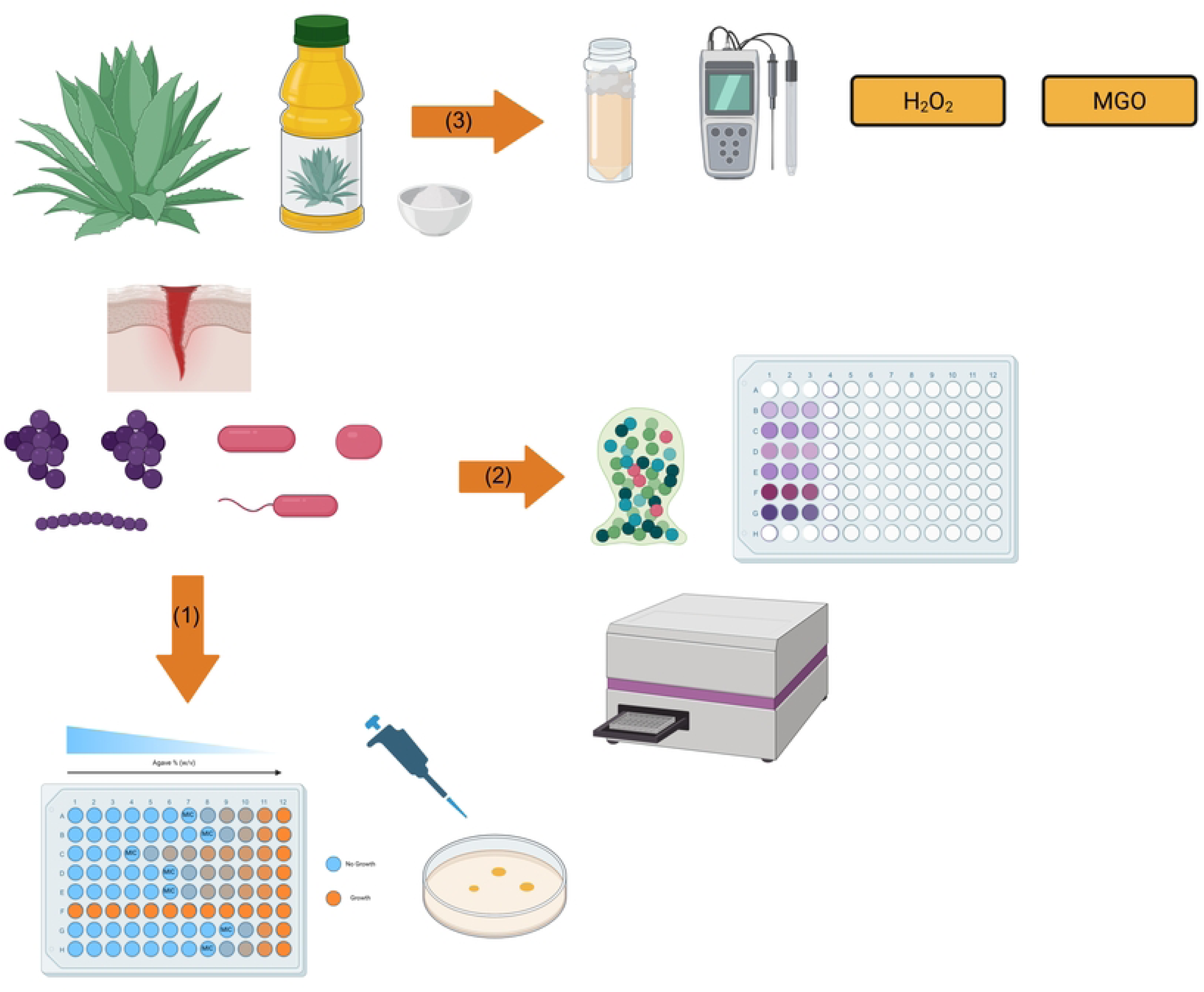

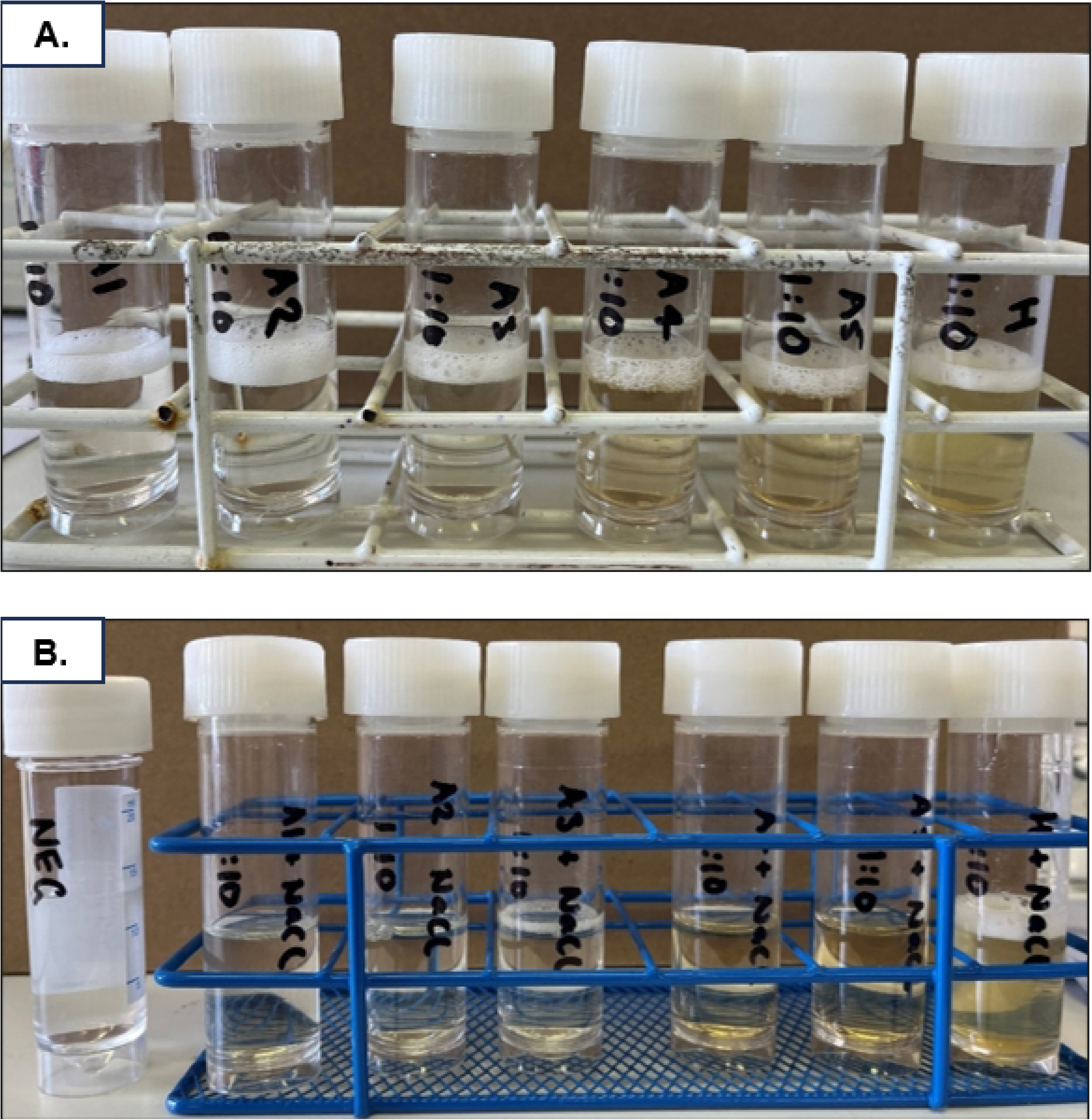

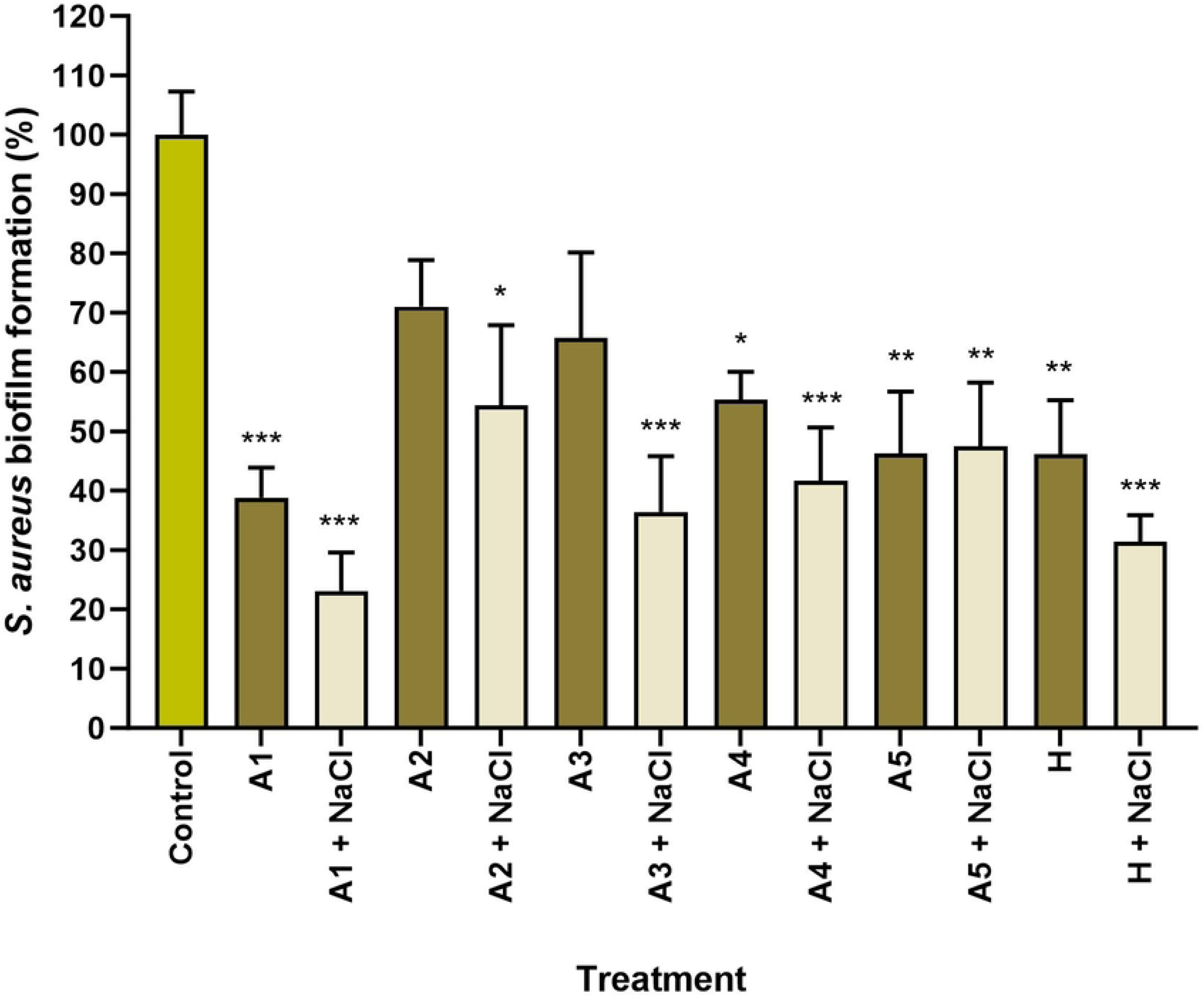

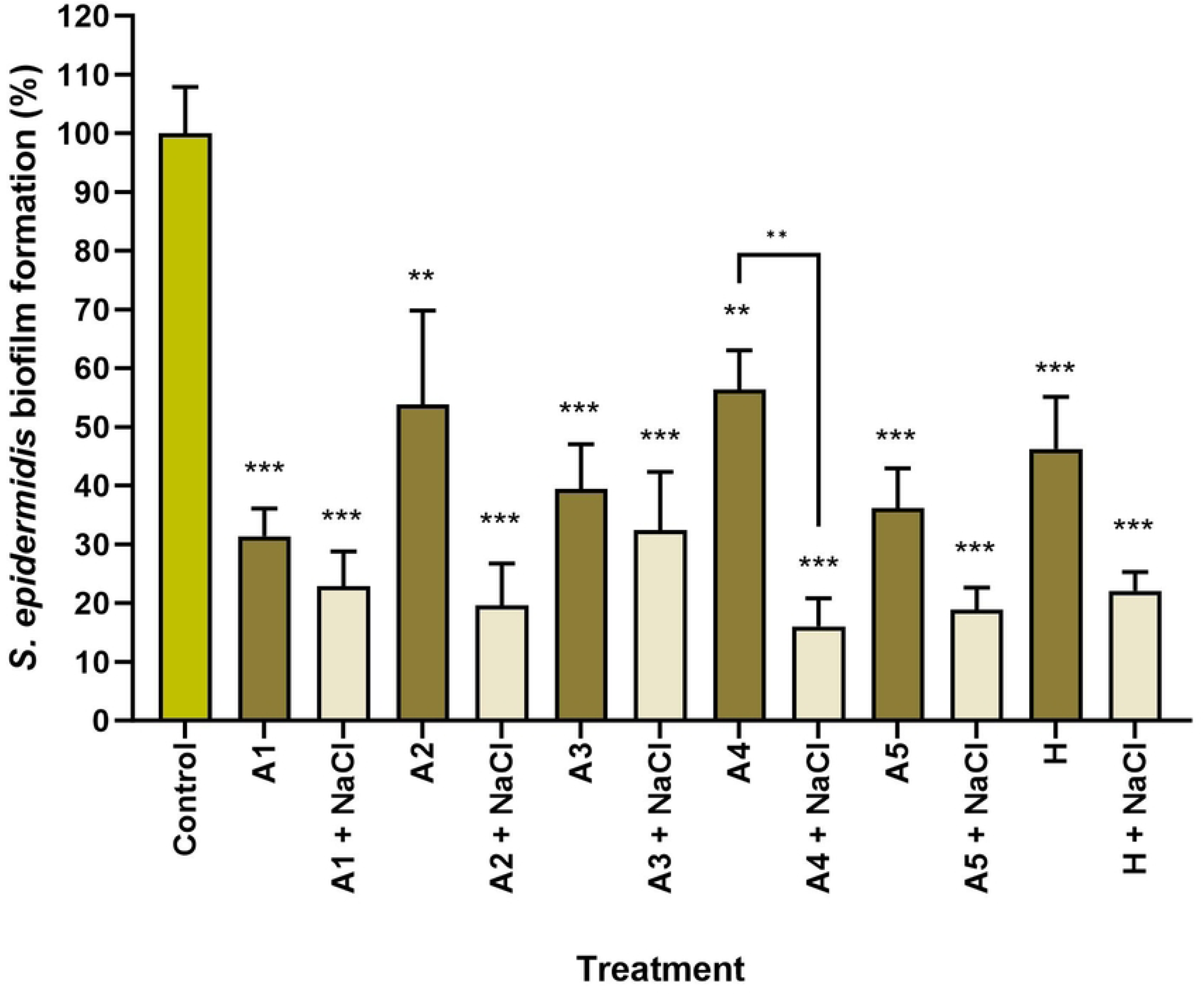

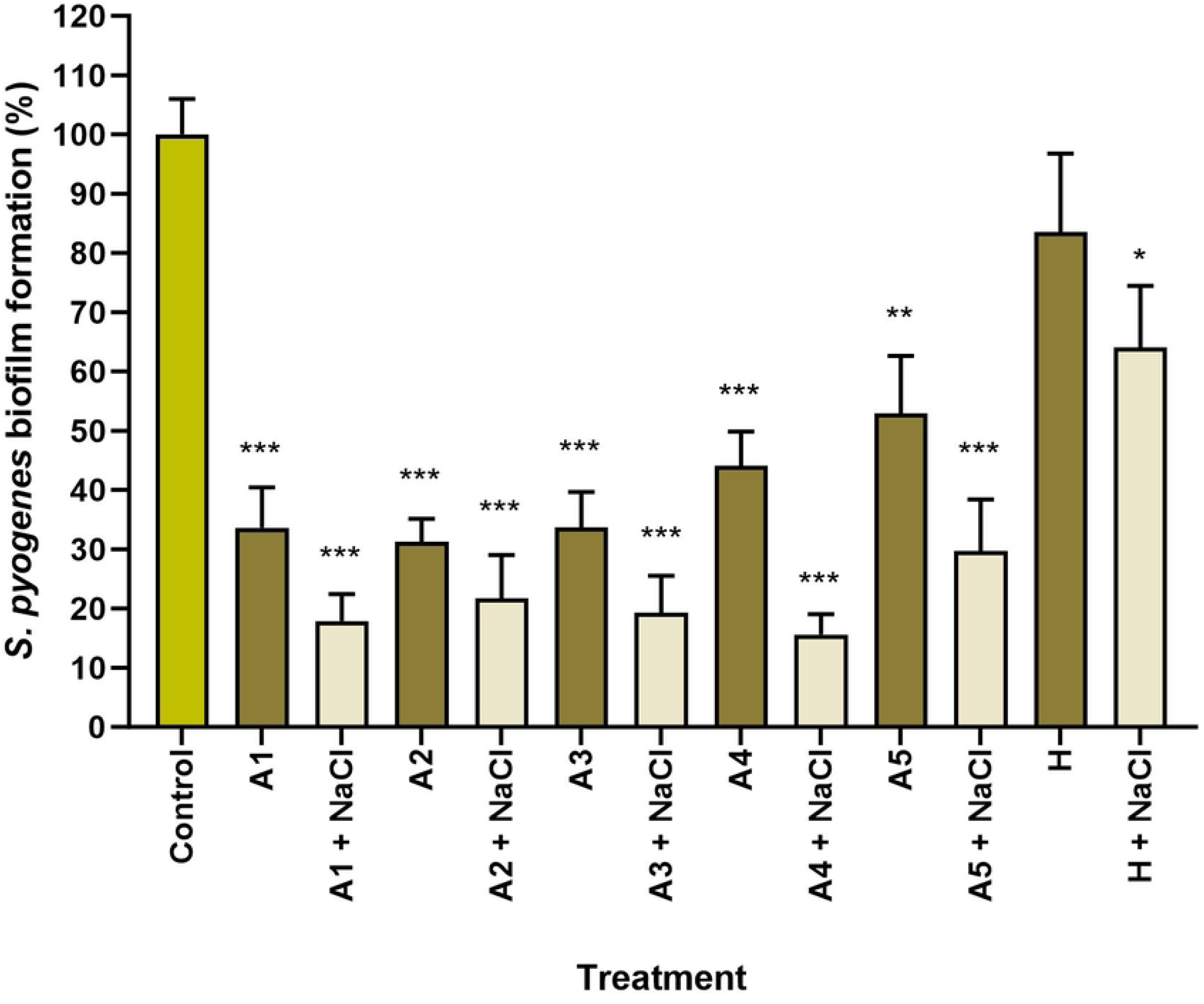

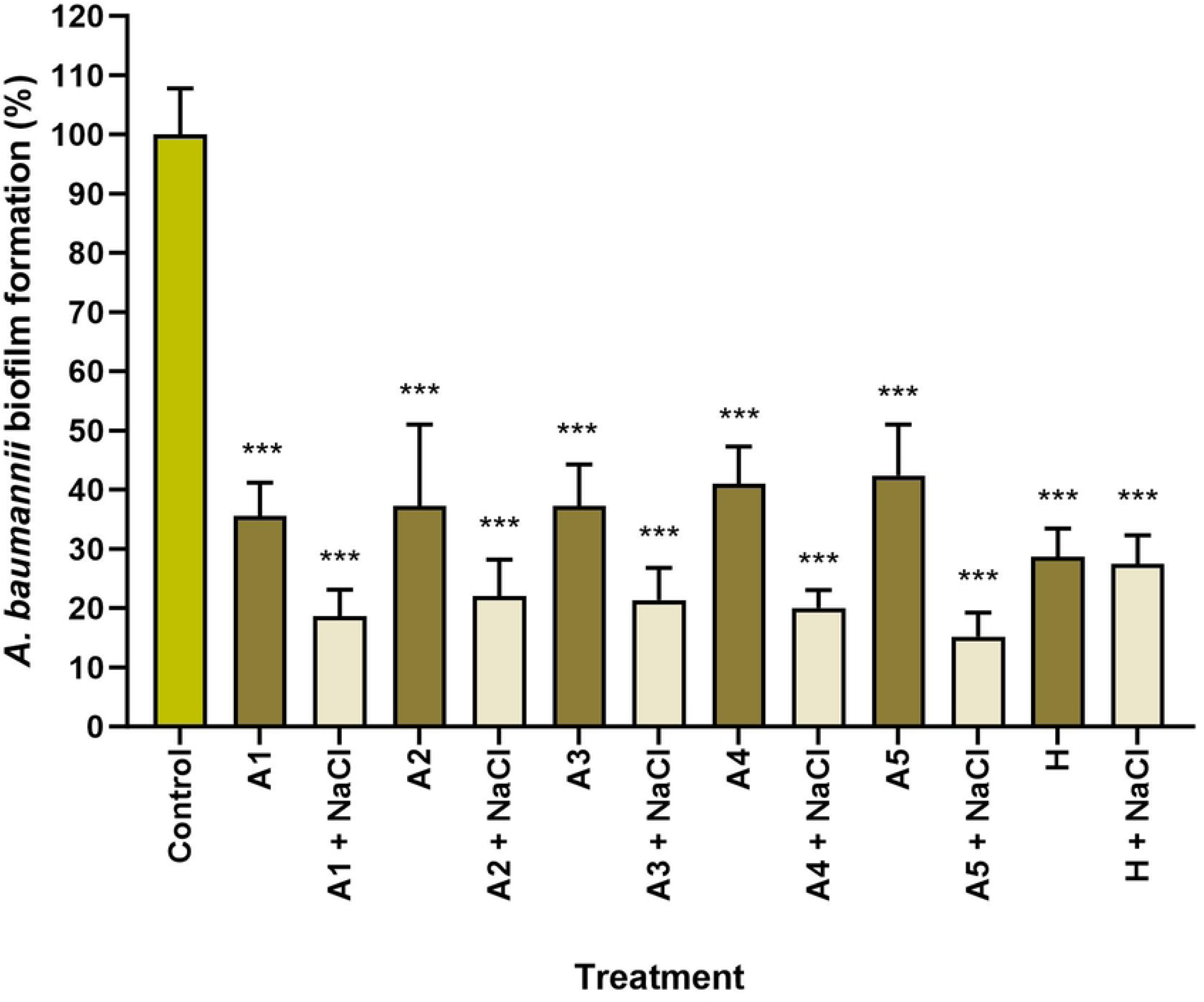

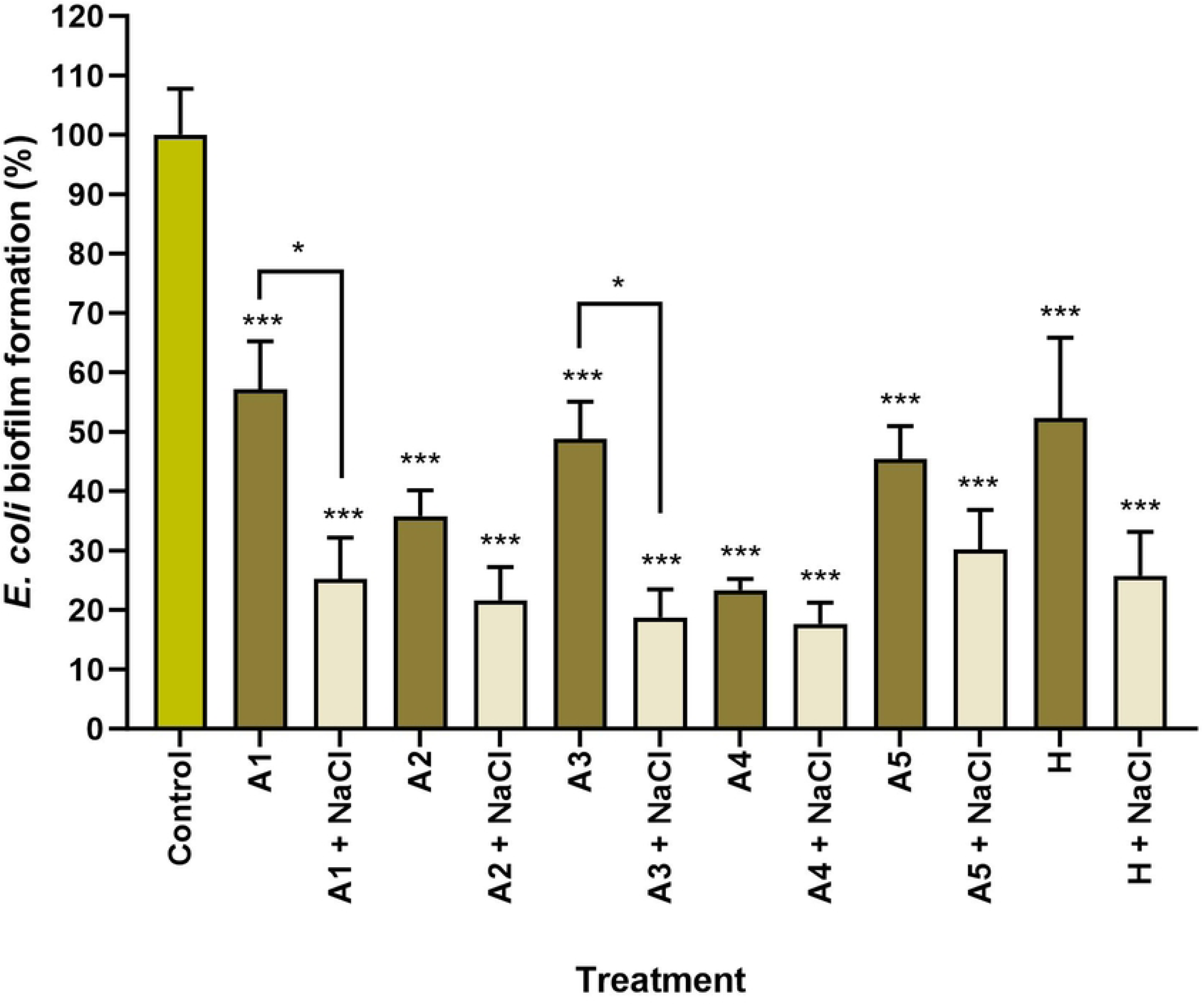

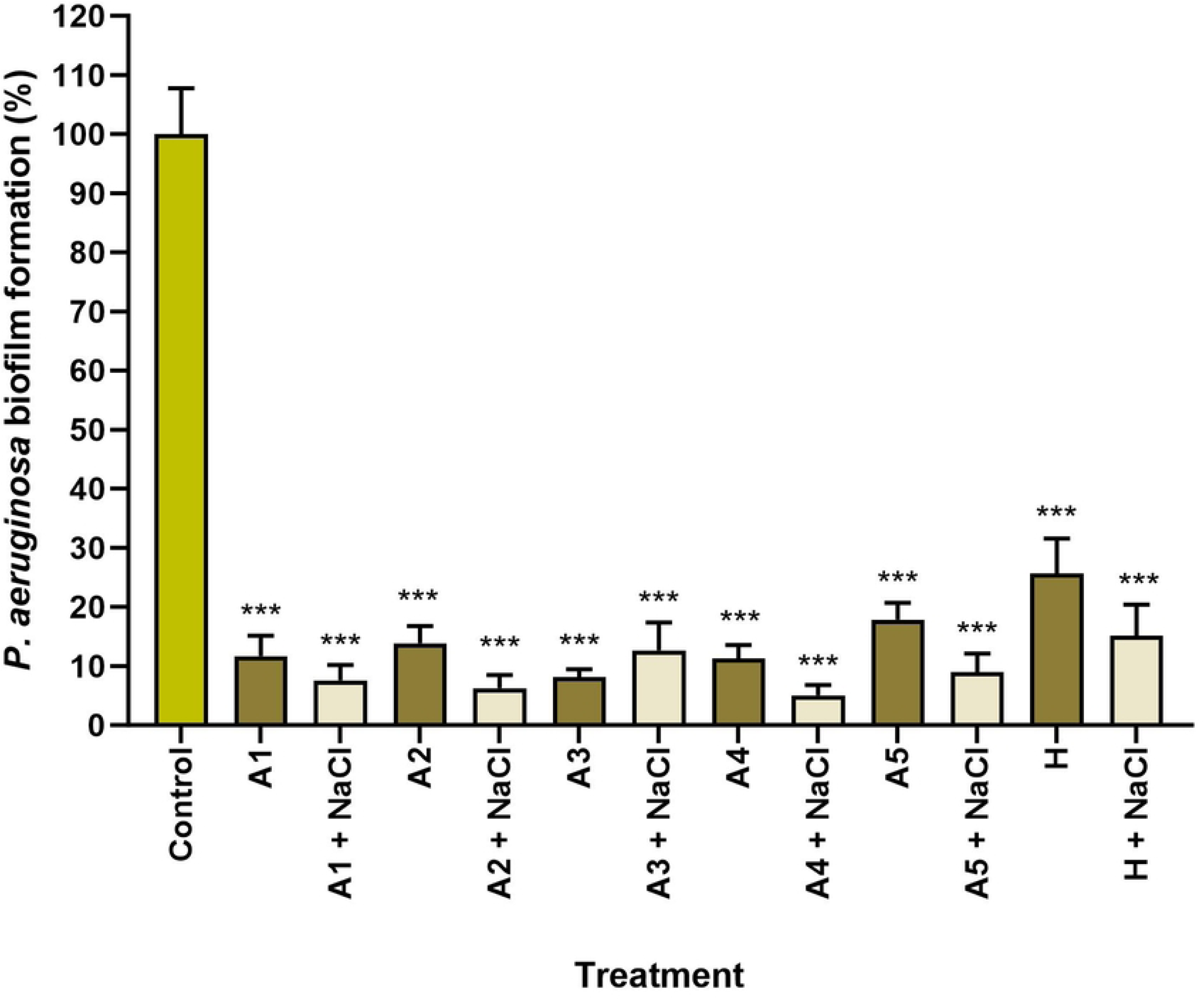

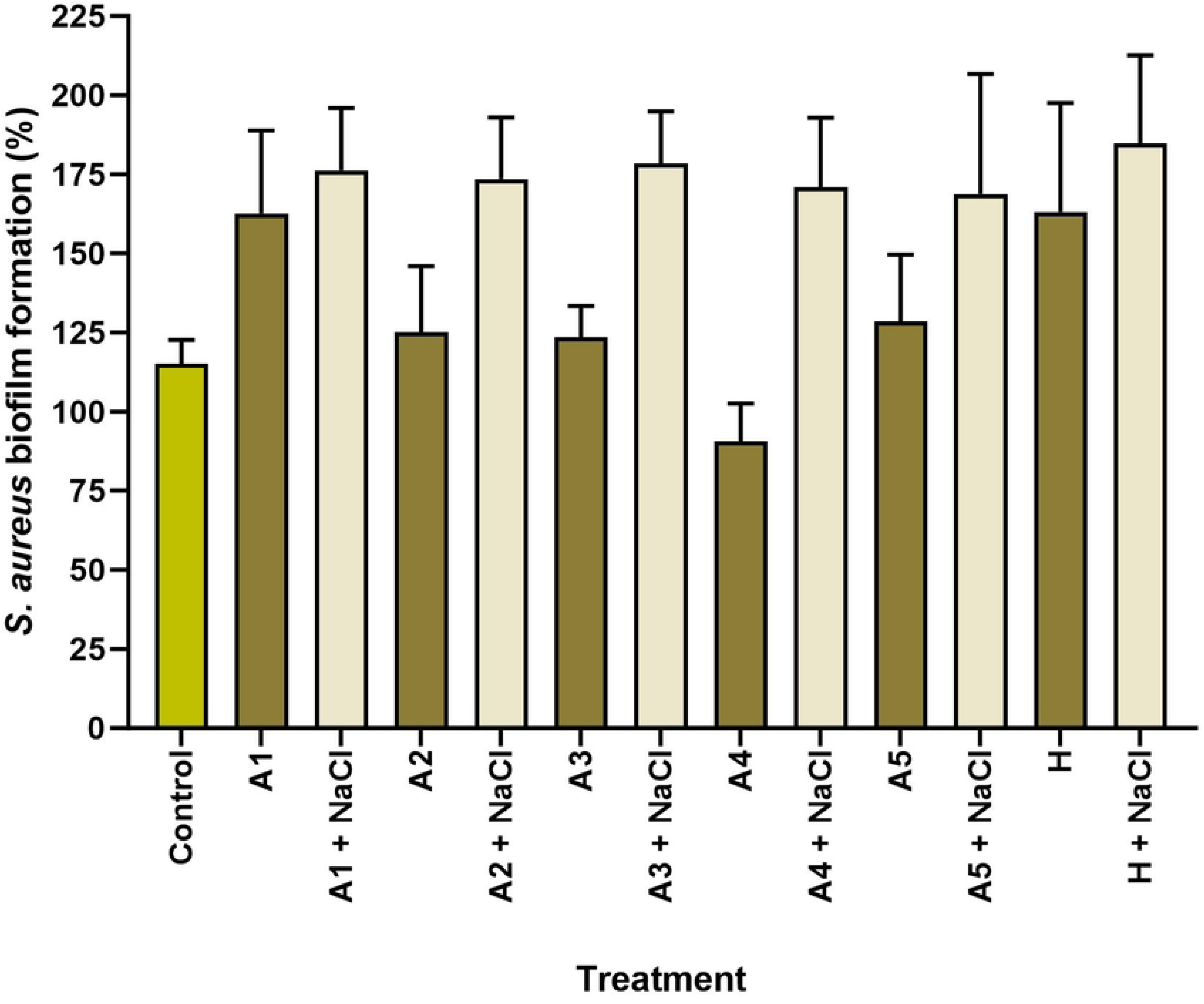

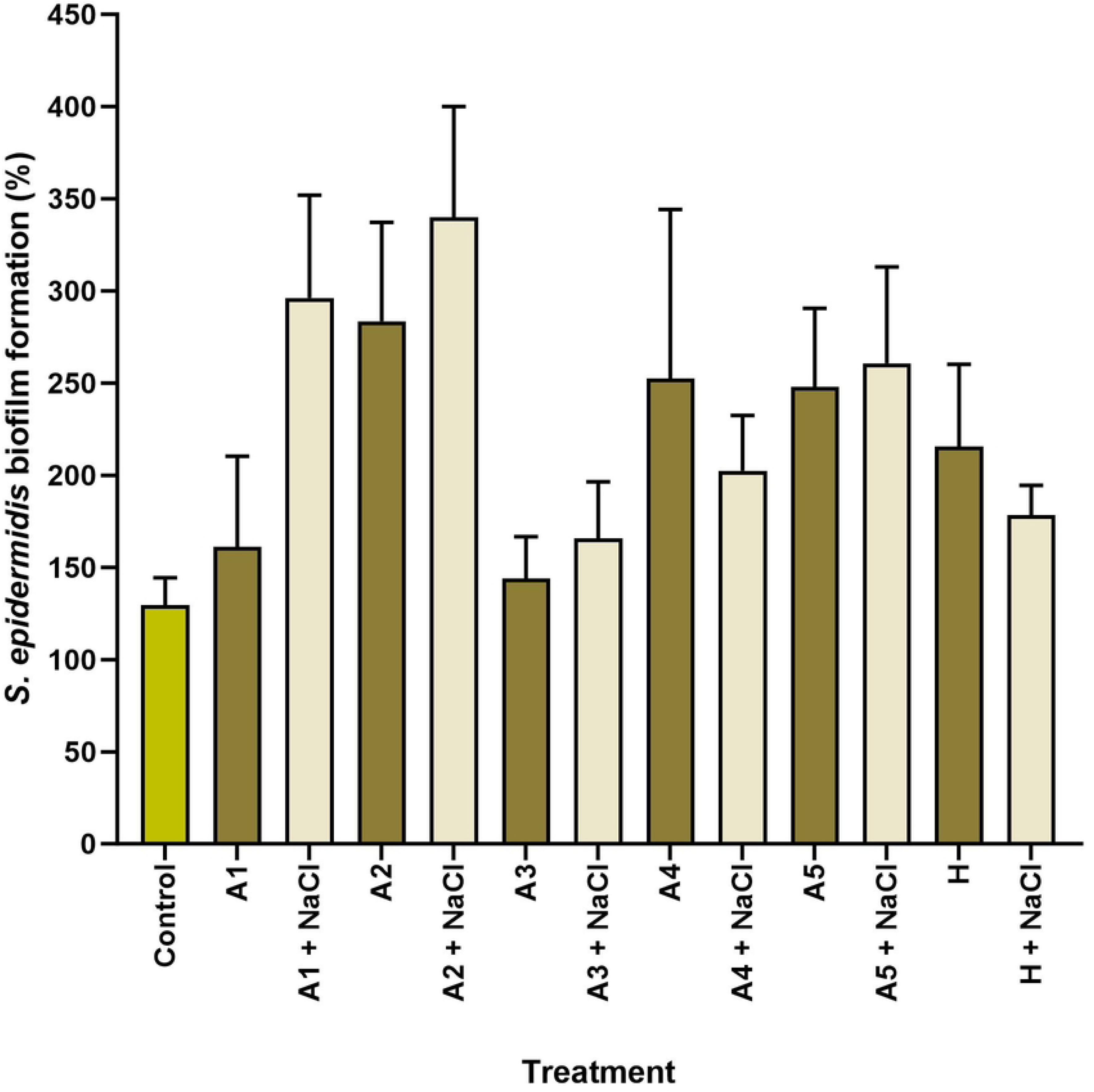

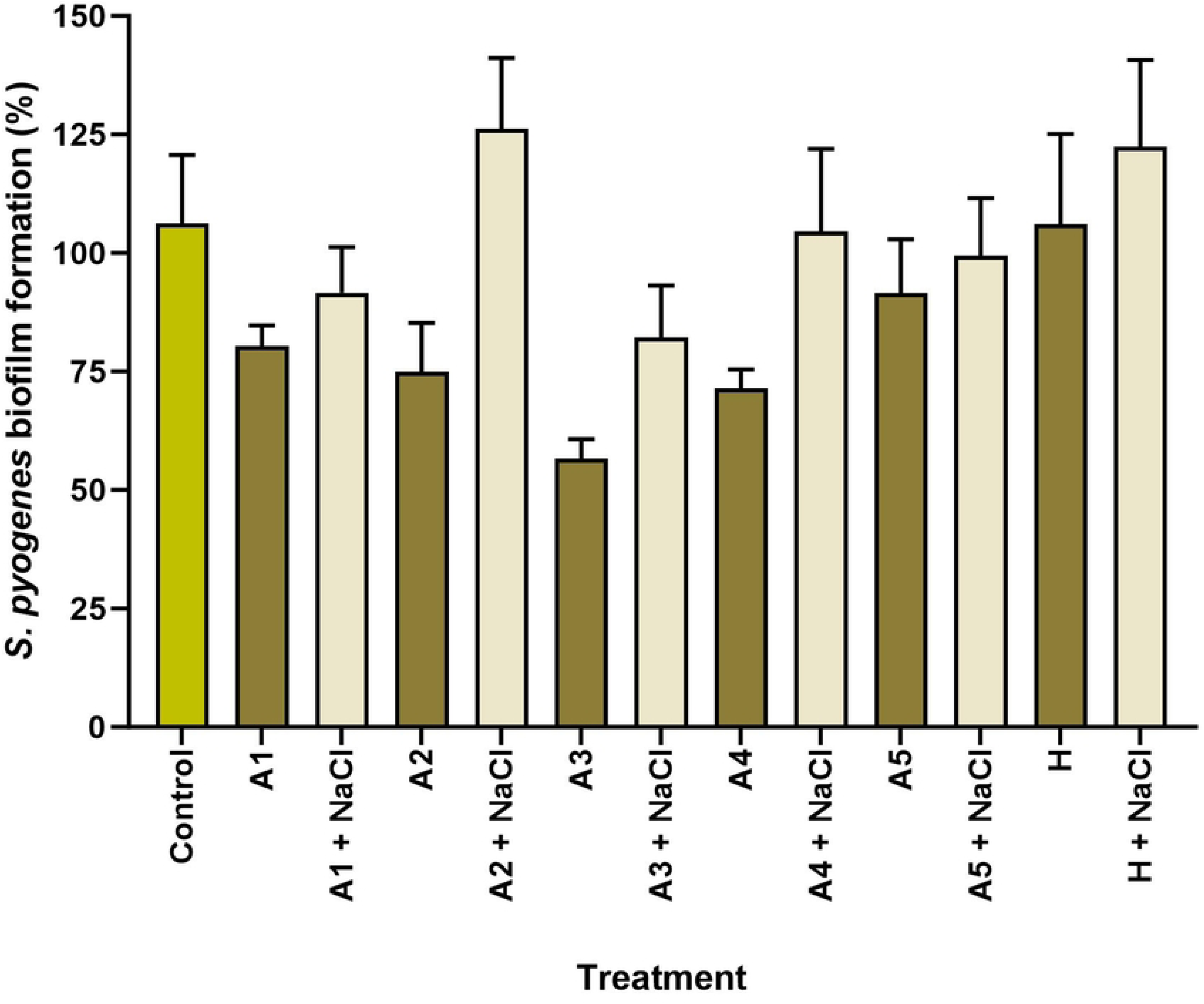

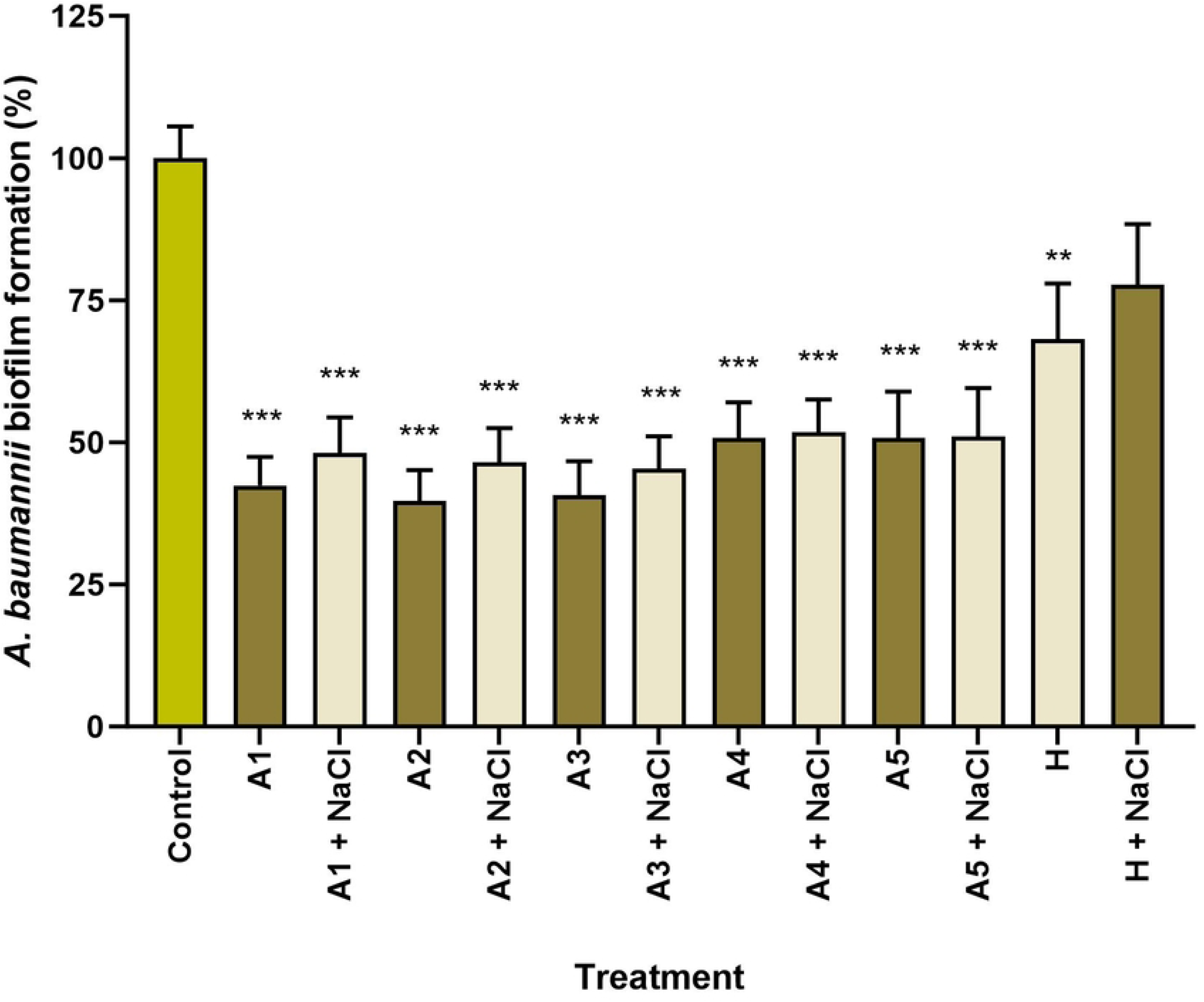

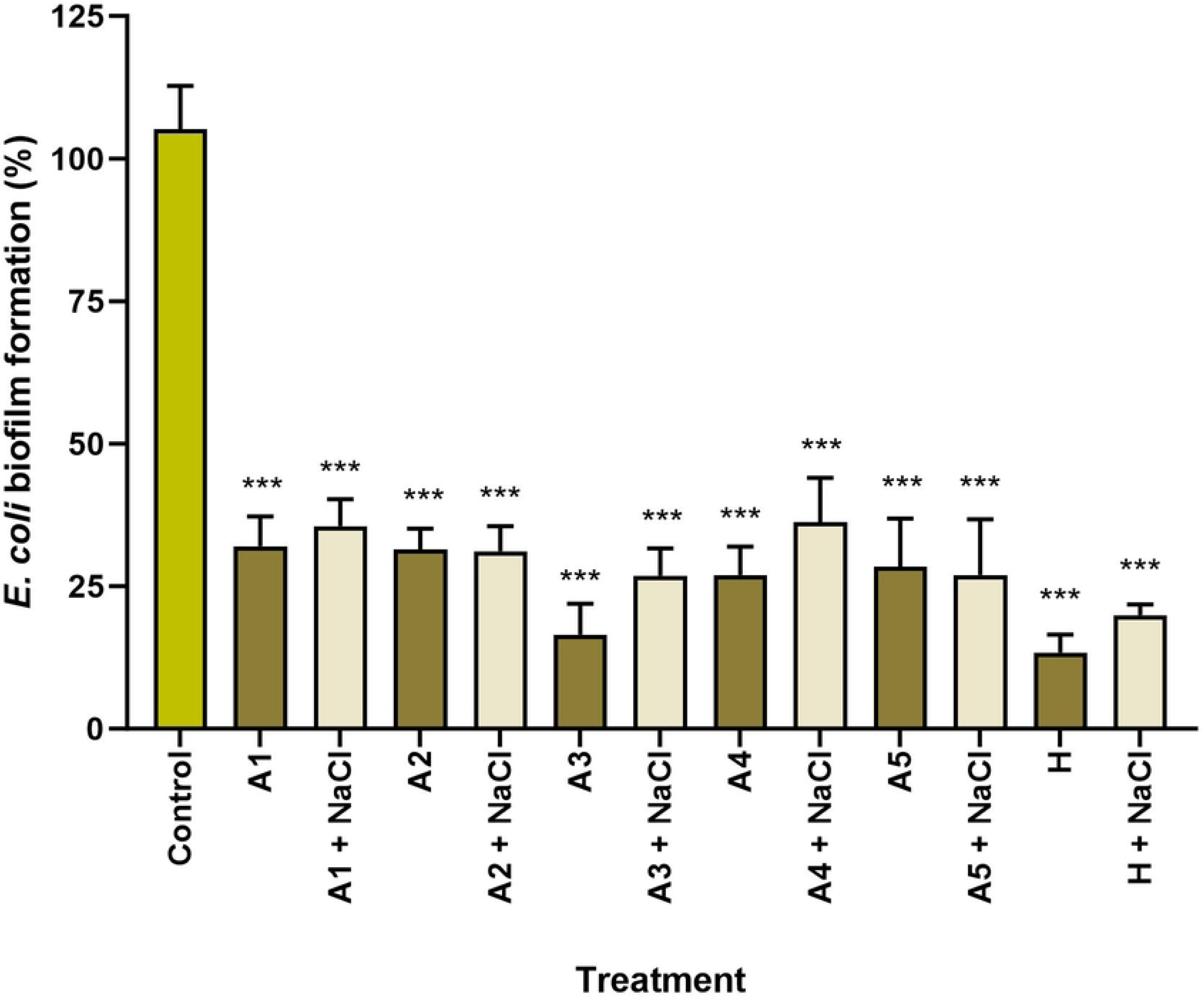

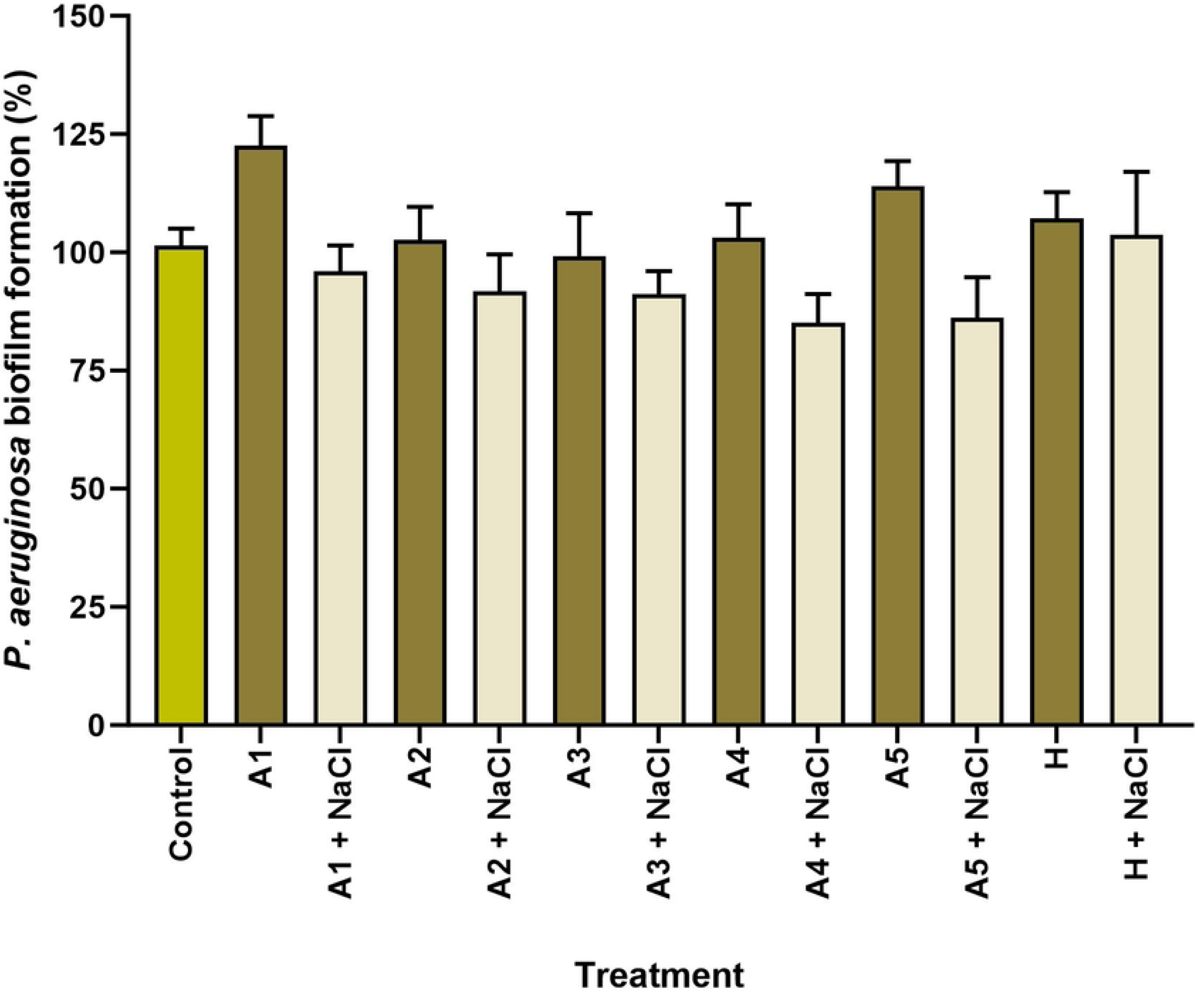

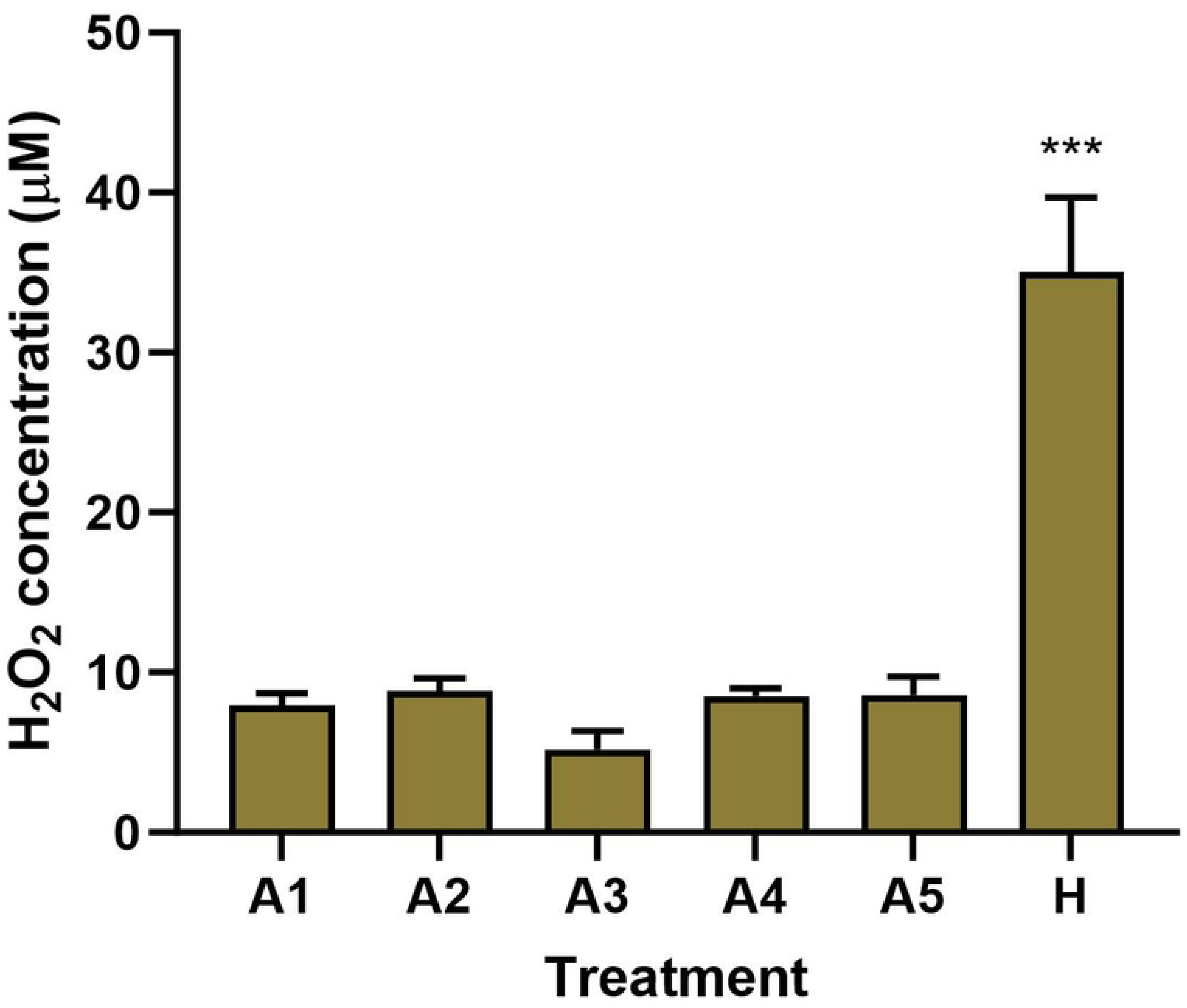

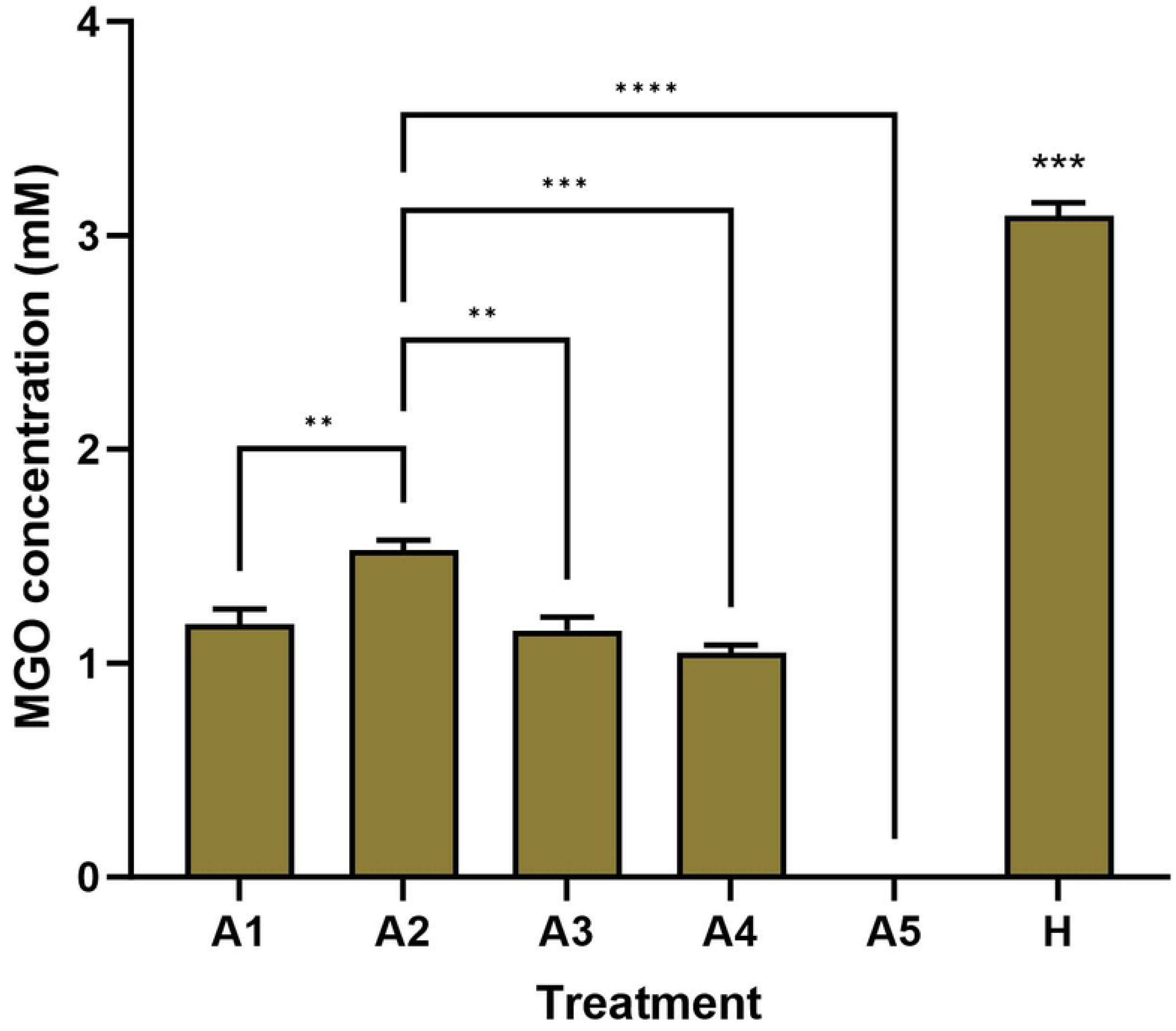

